## Supplementary figures and legends for "Virus and Cell Specific HMGB1 Secretion and Subepithelial Infiltrate Formation in Adenovirus Keratitis"

### Supplemental Figure and Legends:

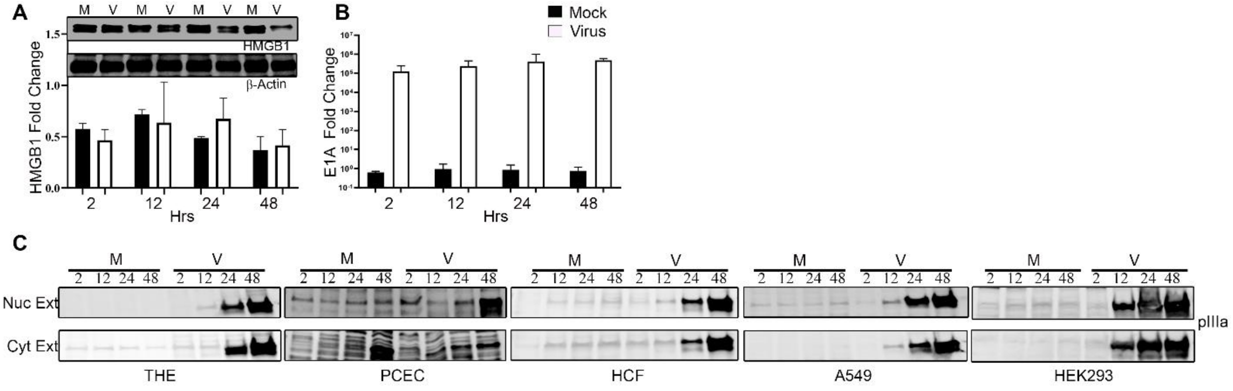


**S1 Fig. Host and viral gene and protein expression in corneal epithelial cells infected with HAdV-D37.** (A) Western blot analysis for HMGB1 expression along with β-actin for load control from whole cell lysates of uninfected (M) or HAdV-D37-infected (V) THE cells for 2, 12, 24, and 48 hpi. qRT-PCR analysis of HMGB1 gene expression for mock and virus infected cells at the same times pi is shown as bar graphs below the Western blot. (B) Bar graph for qRT-PCR of the viral early gene E1A expression, a surrogate marker for viral entry, and normalized to human ACTG gene for quantification. (C) Viral late protein pIIIa expression in cytoplasmic and nuclear extracts prepared from uninfected or HAdV-D37-infected THE, PHCE, HCF, A549 and HEK293 for 2, 12, 24, and 48 hpi show successful infection of all cell types.

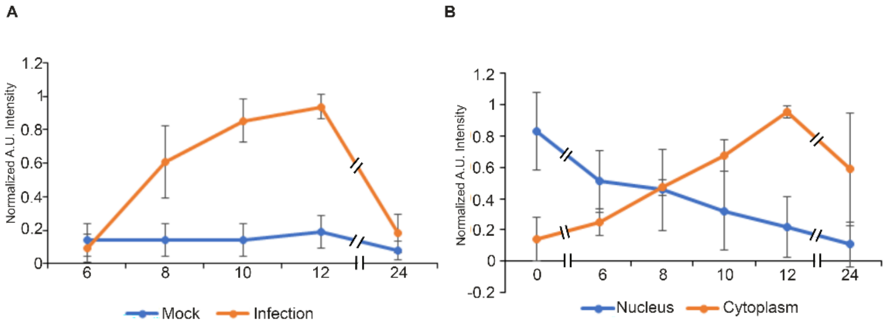

**S2 Fig. Time dependent release of HMGB1 in HAdV-37 infection**. (A) Cytoplasmic HMGB1 localization compared between mock and viral infection at indicated times pi**.** (B) HMGB1 distribution between the nucleus and cytoplasm compared between nucleus and cytoplasm within each group at various time point of infection.

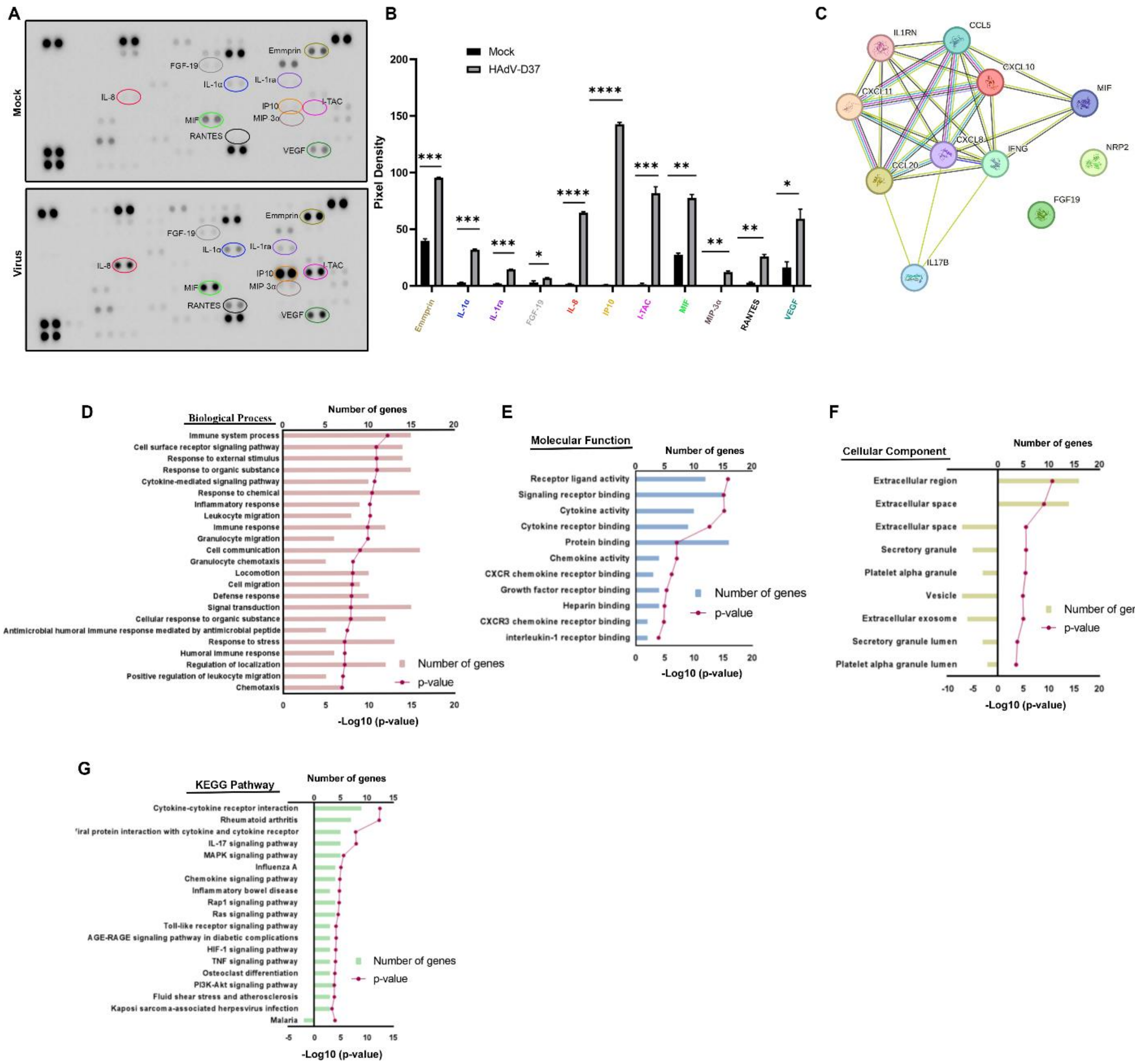


**S3 Fig. Omics analysis of immortalized human corneal epithelial cells infected with HAdV-D37.** (A) Human cytokine array performed on cell supernatant from THE mock treated with dialysis buffer (upper panel) or HAdV-D37 infected (lower panel) at 24 h. (B) Upregulated proteins compared graphically using ImageJ quantification, and compared to same protein in mock infection. (C) The physical and functional associations among the upregulated proinflammatory mediators were assessed using the STRING tool. The interaction among the query proteins represents the network with 10 nodes and 21 edges of protein-protein interaction (PPI). (D-G) Gene ontology (GO) showing top biological process, molecular function, KEGG pathway, and cellular components. Bar length represent the number of genes and dotted line represents -log10 adjusted p value for significantly enriched pathways.

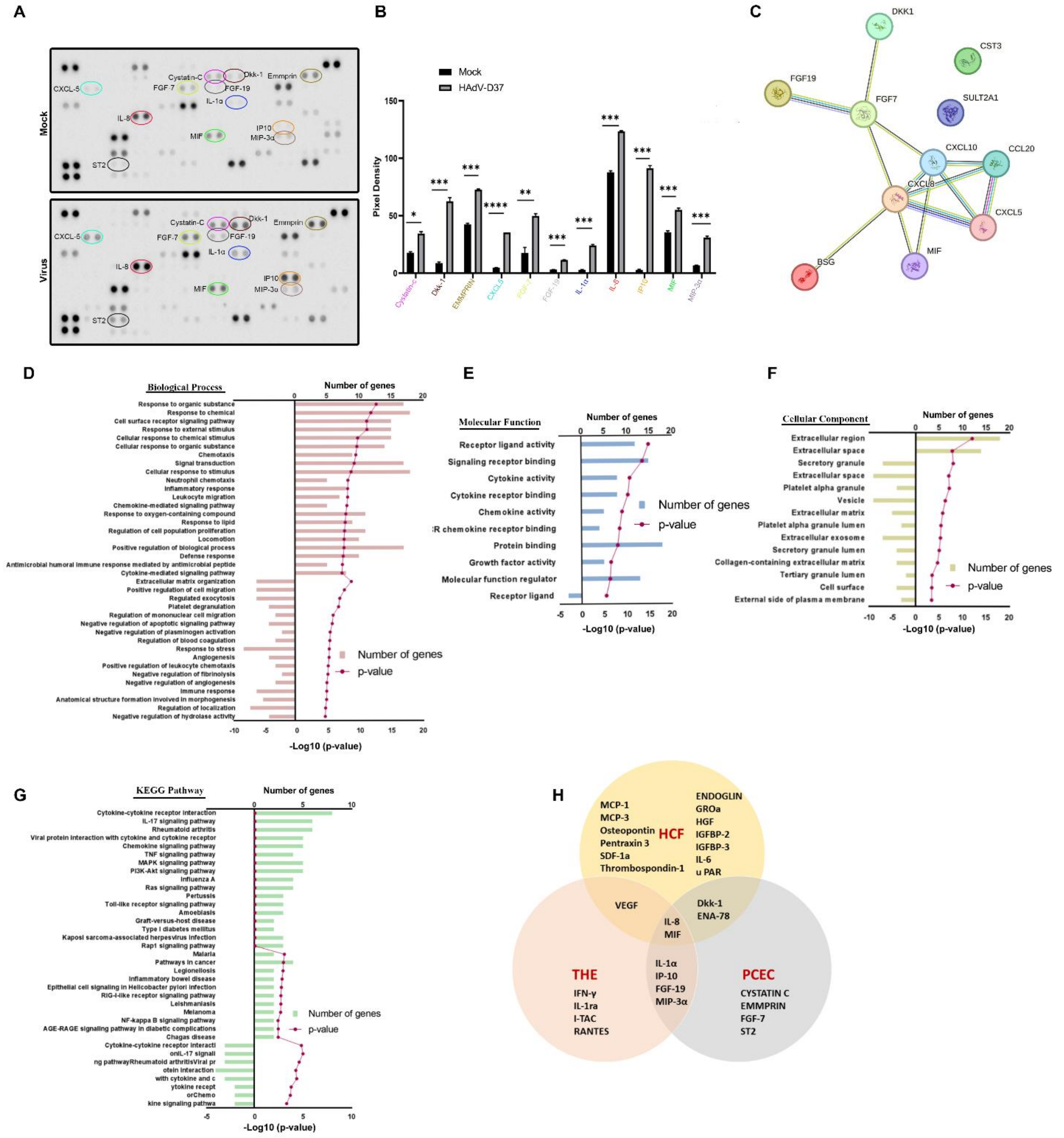


**S4 Fig. Omics analysis of primary human corneal epithelial cells infected with HAdV-D37.** (A) Human cytokine array performed on cell supernatants of, PCEC mock infected (upper panel) or infected with HAdV-D37 (lower panel) at 24 hpi. (B) Upregulated proteins compared graphically using ImageJ quantification, and compared to the corresponding protein in mock infection. (C) The physical and functional associations among the upregulated proinflammatory mediators were assessed using the STRING tool. The interaction among the query proteins represents the network with 11 nodes and 11 edges of protein-protein interaction (PPI). (D-G) Gene ontology (GO) showing top biological process, molecular function, KEGG pathway, and cellular components. Bar lengths represent the number of genes and dotted lines represents -log10 adjusted p values for significantly enriched pathways. (H) Venn diagram showing the upregulated, overlapping cytokine expression between HCF, THE, and PCEC.

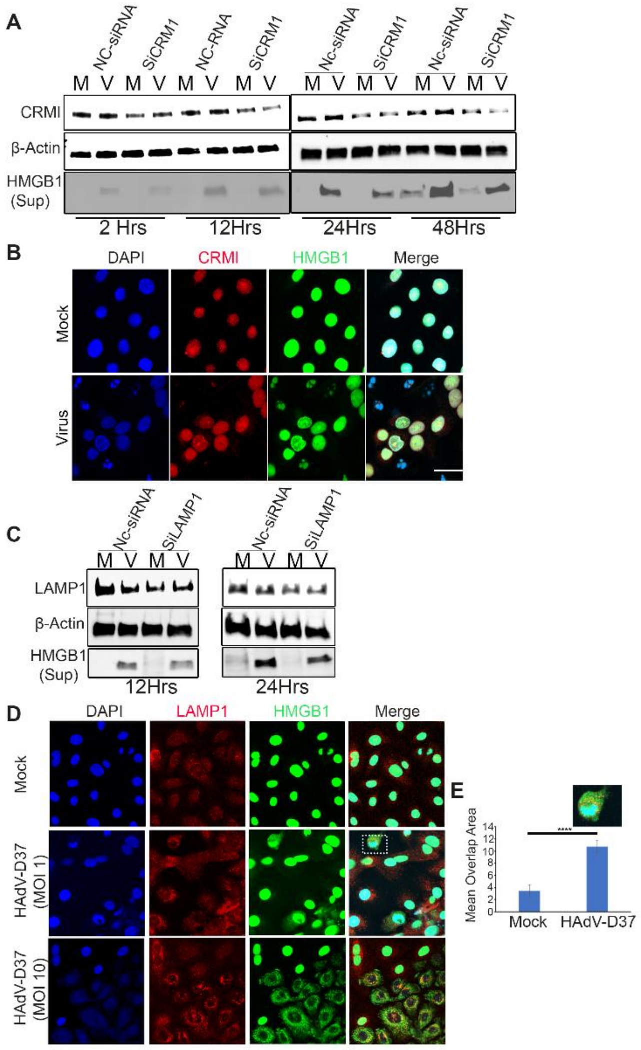


**S5 Fig. Role of CRM1 and LAMP1 in HMGB1 trafficking.** (A) THE cells treated with NC-siRNA or CRM1-siRNA, and HAdV-D37 infected at a MOI of 5 for 2, 12, 24, and 48 h. Cell-free supernatants and total cell lysates were prepared for Western blot to measure HMGB1 and CRM1 knockdown, respectively. β-actin was used as an internal loading control (*n=3*). (B) Immunofluorescence for DAPI nuclear (blue), CRM1 (red), and HMGB1 (green) in mock treated and HAdV-D37 infected cells at an MOI of 5 at 12 hpi. Colocalization between HMGB1 and CRM1 (yellow nucleus) was seen only in virus infected cells. Scale bar =10 μm. (C) THE cells were treated with NC-siRNA and LAMP1-siRNA, and HAdV-D37 infected at an MOI of 5 for 12 and 24 h. Cell-free supernatants and total cell lysates were prepared to measure HMGB1 and LAMP1 knockdown respectively by Western blot. β-actin used as an internal loading control (*n=3*). (D) Immunofluorescence showing nuclei (blue), LAMP1 (red), and HMGB1 (green) in mock infected and HAdV-D37 infected cells (MOI of 1 and 10, at 12 hpi). Scale bar = 10 μm. (*n=5*). (E) HCM analysis of THE cells infected with HAdV-D37, and immunostained for HMGB1 and LAMP1 and analyzed for colocalization. Data shown as the mean ± SD (*n= 3*). ANOVA with Tukey's post-hoc test was performed. For HCM, >5000 cells were counted per well, with a minimum number of 3 valid wells (*n=3*).

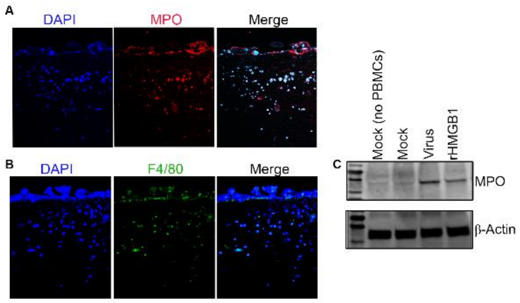


**S6 Fig. MPO and F4/80 expression in infected 3D corneal constructs.** (A) Sections from 3D corneal constructs were stained with DAPI, and immunostained for MPO expression**.** (B) Sections stained with DAPI and immunostained for F4/80 expression. Figures are representative of images from three 3D corneal constructs. (C) Western blot analysis showing expression of MPO in constructs infected with HAdV-D37 or treated with rHMGB1.
